# Distance-AF Multimer: A Multi-Step Approach for Protein Complex Structure Modeling with User-Defined Distance Constraints

**DOI:** 10.64898/2026.10.14.759232

**Authors:** Kai Ling, Yuanyuan Zhang, Zicong Zhang, Tsukasa Nakamura, Daisuke Kihara

## Abstract

The three-dimensional structure of proteins provides essential insight into the molecular mechanisms underlying their function. Many biological activities are mediated through protein interactions, making accurate structural determination of protein complexes important. Despite significant advances in computational protein structure prediction, there are still many cases where protein complexes are not correctly assembled. In particular, there are cases where experimental data, such as distances between residue pairs in interacting proteins are available, yet existing prediction methods fail to produce models consistent with these data. Here, we present Distance-AF Multimer, a method that guides and refines AlphaFold-Multimer docking predictions using user-provided residue pair distance information. This approach extends our previously successful data-guided single-protein modeling method, Distance-AF. Distance-AF Multimer operates in a two-step process, starting from multiple initial conformations and iteratively refining them to satisfy the provided distance constraints. We evaluated Distance-AF Multimer on protein-complex benchmarks spanning different chain counts and docking errors, demonstrating competitive or superior performance compared to other existing deep-learning-based structure prediction methods. In addition to dimer cases, we also demonstrated the potential of Distance-AF Multimer for large-scale protein assemblies containing multiple chains, including those with more than 5 chains.

**Significance Statement:** Proteins often function as complexes, but predicting how their components fit together remains difficult. Distance-AF Multimer uses distance constraints between residues to improve the arrangement of protein chains in predicted structures. This approach offers a practical way to combine computational predictions with available structural information, helping researchers study protein interactions, peptide binding, and larger assemblies when detailed experimental structures are unavailable.

## Introduction

Multi-chain protein complexes mediate the majority of cellular functions, ranging from signal transduction to catalysis and regulation (Lu et al., 2025). While experimental techniques such as X-ray crystallography and cryo-electron microscopy (cryo-EM) provide invaluable structural insights, these approaches often require substantial time and resources. To complement experimental methods, computational approaches for protein complex structure prediction have been extensively studied (Aderinwale et al., 2020).

In the context of single chain protein structure prediction, recent deep learning breakthroughs, most notably AlphaFold2 (Jumper et al., 2021), have achieved remarkable success and have been widely discussed in recent years. However, predicting protein complex structures remains challenging in many practical scenarios. This difficulty arises from several factors that increase the problem complexity, including the larger size of complexes, conformational flexibility at protein-protein interfaces that can lead to structural differences from isolated chains, and weaker evolutionary signals across interacting chains compared to those within single proteins. As a result, current methods frequently exhibit failures in accurately predicting interface geometries and often produce substantial domain orientation errors (Kagaya et al., 2026; Lu et al., 2025; Zhang et al., 2026).

To increase the accuracy of modeling, one approach is to integrate experimental data into the protein structure prediction procedure, if available. Many conventional rigid-body protein docking methods, such as LZerD (Christoffer et al., 2021; Harini et al., 2023; Venkatraman et al., 2009), HADDOCK (de Vries et al., 2010) and ClusPro (Kozakov et al., 2017) implemented experimental restraints, i.e. residue pair distances and interface residues, into structure prediction workflows. Methods based on RosettaDock (Lyskov and Gray, 2008) integrated crosslinking scoring terms into their scoring functions for model selection. AlphaLink2 (Stahl et al., 2024) advanced this approach by embedding crosslinking mass spectrometry information into AlphaFold2’s pair representation (Jumper et al., 2021), with an extension for protein complex prediction. Recent diffusion model-based multi-chain structure modeling methods, such as Boltz-2 (Passaro et al., 2025), Chai-1 (Boitreaud et al., 2024), and Protenix (Chen et al., 2025), have incorporated constraint features during training. During inference, these models can leverage experimental data that provides residue pair distance information, such as crosslinking mass spectrometry or epitope mapping, to improve prediction accuracy. However, existing approaches remain constrained by several limitations: they require substantial numbers of restraints to achieve meaningful improvements and exhibit suboptimal satisfaction of imposed constraints. Moreover, deep learning-based methods require extensive retraining on large-scale experimental datasets to incorporate constraint-handling capabilities, limiting their adaptability and accessibility. Distance-AF (Zhang et al., 2025a), which we developed for a single-protein structure modeling, offers a fundamentally different paradigm through post-hoc constraint-guided refinement of AlphaFold2 predictions, requiring neither model retraining nor large annotated datasets. Here, we present Distance-AF Multimer (Distance-AFM), extending this training-free constraint-based approach to multi-chain protein complexes where the challenges are fundamentally different and more complex than single-chain prediction.

Distance-AFM addresses limitations that cannot be effectively resolved by existing constraint-based docking methods. Unlike single-chain predictions, for which AlphaFold2 typically provides reasonable initial structures, multimeric complex predictions often suffer from severely incorrect inter-protein interactions, including large rotational errors that cannot be corrected by standard distance-based refinement. To address this challenge, we employ a two-step strategy to correct inconsistencies in predicted docking models, primarily those generated by AlphaFold-Multimer (AFM) (Evans et al., 2022), using known distance constraints. Distance-AFM first performs a pre-estimation step to generate diverse starting conformations, followed by the incorporation of distance constraints to guide optimization toward geometrically consistent structures. This strategy overcomes the limitation of directly applying distance-driven optimization to AlphaFold-Multimer predictions, which frequently fails due to severe initial pose errors.

Distance-AFM demonstrates robust performance across diverse protein complex types and scales. For challenging targets where AFM produces severely misfolded initial predictions, our method successfully recovers near-native structures through constraint-guided refinement. Benchmarking against state-of-the-art models including Chai-1 (Boitreaud et al., 2024) and Protenix (Chen et al., 2025) on selected protein-complex benchmarks shows that Distance-AFM achieves competitive or superior accuracy while maintaining the unique capability to generate multiple conformationally distinct states. The method scales effectively to large assemblies, successfully modeling systems with up to 10 chains, a capability particularly valuable for studying biological machines such as viral capsids, ribosomes, and multi-enzyme complexes where conformational flexibility plays critical functional roles.

## Results

### Three-Phase Architecture of Distance-AF Multimer

Distance-AFM extends the Distance-AF (Zhang et al., 2025a) framework, originally developed for single-chain proteins, to multi-chain protein complexes. The workflow of Distance-AFM is shown in **Fig. 1**. It introduces a coarse tuning phase to the original Distance-AF, in which multiple starting structures are generated, each of which is optimized using the provided distance constraints. The inputs of Distance-AFM consist of the protein sequences to be docked and a list of inter-chain amino acid residue pairs with associated distance information.

**Figure 1.**
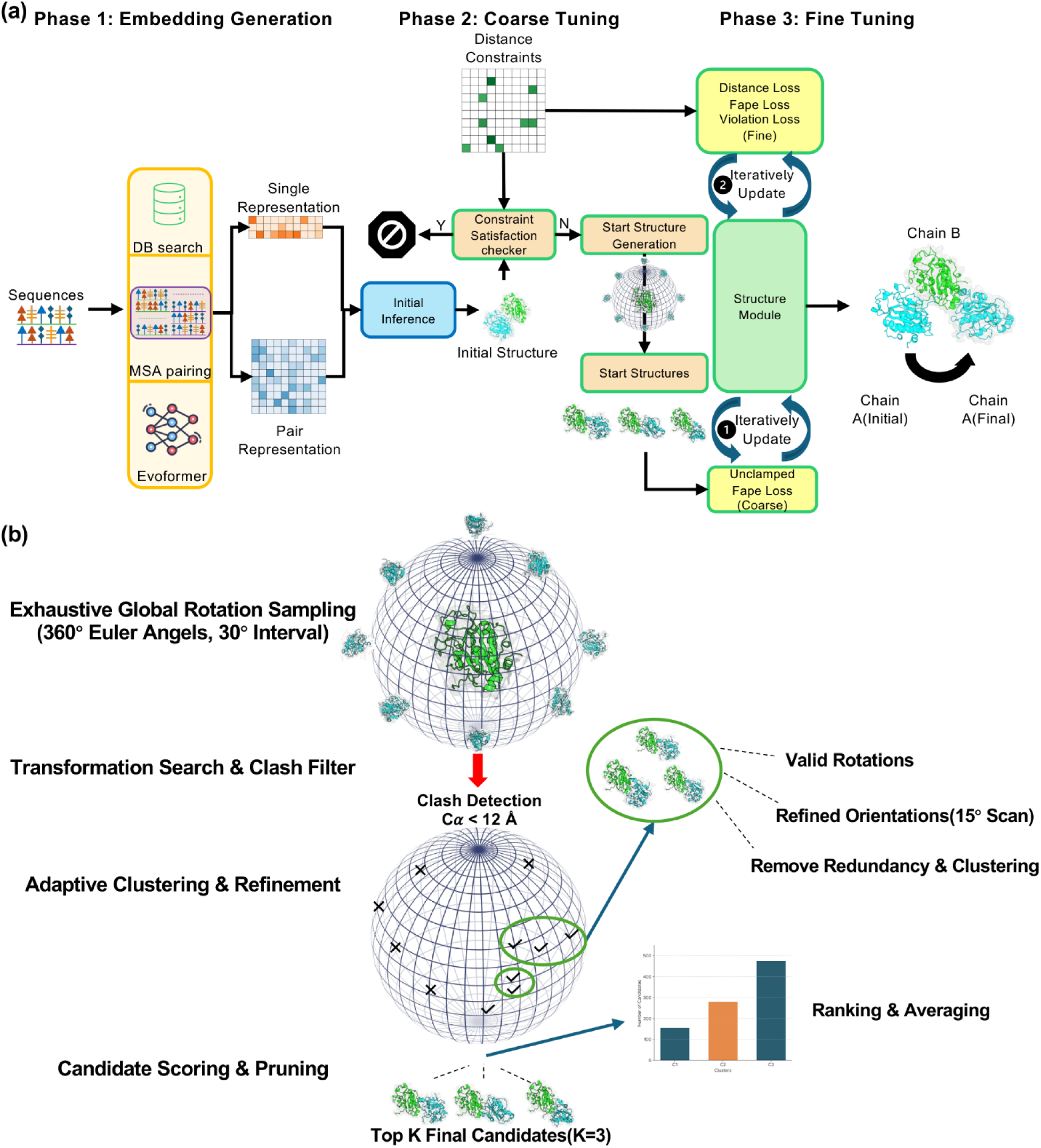
Overall framework of Distance-AF Multimer. **(a)** Overview of the three-phase pipeline for constraint-guided multimer structure prediction. Phase 1 (Embedding Generation): Sequences are processed through homology search, MSA pairing, and Evoformer encoding to generate single and pair representations. Phase 2 (Coarse Tuning): Distance constraints are integrated through a two-step preprocessing protocol, which explores docking poses at different orientations while satisfying a selected user-provided distance constraint. Then, valid orientations are clustered by adjacency in Euler-angle space; clusters are ranked by cluster size (number of valid poses), and the top K clusters (K = 3) are retained. Each selected cluster is summarized by averaging its Euler angles (α, β, γ) to obtain a representatives rotation used to initialize the next refinement step. Phase 3 (Fine Tuning): The Structure Module iteratively optimizes the structure using FAPE Warm-up Loss, Distance Loss, and FAPE Loss through iterative weight updates. Green boxes indicate standard AlphaFold-Multimer components, yellow boxes represent constraint-specific modifications. The preprocessing phase (Phase 2) enables correction of large rotation errors that cannot be addressed through standard fine-tuning alone. **(b)** Overview of Phase 2, Coarse-Tuning starting structures. At each iteration of orientation generation, one chain is selected as the primary moving unit while the remaining chains are fixed. Exhaustive global rotations are first sampled using discretized Euler angles, followed by clash filtering based on Cα distance thresholds to remove sterically infeasible conformations. The remaining valid orientations are clustered, locally refined with finer angular scans, and used to generate candidate structures for the next assembly stage.

The procedure starts from running AFM. In this first phase, Distance-AFM generates single and pair representations for each chain in the complex, alongside inter-chain pair representations, following the AFM protocol (Evans et al., 2022). This involves paired MSA (Morcos et al., 2011) construction for the protein chains by searching UniRef90 (UniProt, 2025), MGnify (Richardson et al., 2023), UniClust30 (Mirdita et al., 2017), BFD (Steinegger and Soding, 2018), and inter-chain pairing is performed by grouping sequences by species using labels from the UniProt (UniProt, 2025) idmapping database (Evans et al., 2022). Structural templates may be identified for individual chains using the PDB Seqres database. In a practical scenario, the AFM complex model is then evaluated against the user-defined distance constraints. Models that do not satisfy these constraints are passed to Phase 2.

This second phase introduces a two-step coarse-to-fine tuning preprocessing, which generates a set of starting structures for distance-based structure optimization (**Fig. 1b**). Since distances between specified residue pairs are adjusted using a gradient-based approach that mainly induces translational movements of proteins, it becomes challenging for Distance-AFM when the starting structure contains large rotational errors. Starting from an initial AFM prediction, we construct candidate conformations by treating each chain (or chain group) as a rigid body and explicitly searching relative transformations between chains under the provided inter-chain distance constraints. More concretely, we first compute a constraint-consistent translation of the chains and then perform an exhaustive rotational sweep with an interval of 30 degrees for three Euler angles (α, β, γ) in the Euler-angle space. Typically, 10^3^–10^4^ docking conformations are generated for a complex of two proteins. Candidate poses are filtered by steric feasibility using a clash criterion (inter-chain Cα–Cα distance < 12 Å), and the remaining valid orientations are clustered based on adjacency in the Euler-angle space. Neighboring grid orientations differ by one 30° step in exactly one of three Euler angles (α, β, γ), while the other two angles remain unchanged. Clusters are constructed by grouping mutually adjacent rotations, thereby removing near-duplicate orientations arising from the discrete rotational sampling. To further improve cluster continuity, each cluster is locally refined by exploring angular gaps between neighboring rotations using a finer rotational interval of 15°. Additional rotations sampled within these gaps are incorporated into the cluster if they satisfy the steric feasibility criteria, ensuring that contiguous regions of valid orientation space are fully captured. After clustering and local gap refinement, the three largest clusters are selected. For each selected cluster, a representative rotation is computed by taking the mean of the three Euler angles (α, β, γ) over all rotations contained in the cluster. The resulting three averaged rotations are used to generate three geometrically distinct starting conformations, which are then passed to the final fine-tuning phase. The preprocessing algorithm and its implementation details are provided in Supplementary Information 1. This procedure is illustrated in **Fig. 1b**.

In Phase 3, starting from the conformations generated in the previous phase, the structure models are refined by optimizing residue-pair distances according to user-defined distance constraints. These constraints specify distances between selected residue pairs at their Cα atoms. To evaluate satisfaction of the distance constraints, the following loss function is added to the loss functions originally implemented in AF-Multimer:

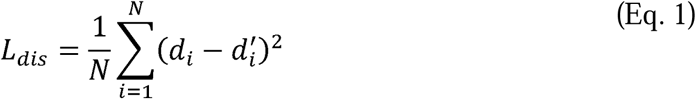

where *N* is the number of distance-constraint pairs; *i* indexes the *i*-th constraint pair; *d_i_* and *d’_i_* are the target and predicted Cα–Cα distances for that pair, respectively.

This distance loss is combined with four loss terms that originate from AF-Multimer: They include a center-of-mass loss (Lcom), which is employed to penalize large-scale translational offsets between chains, and a chain clash violation (*L_vio_*) loss, which prevents non-physical steric overlaps at the interfaces. In addition, we include an angle loss (*L_angle_*), inherited from AlphaFold-Multimer, which penalizes deviations in predicted backbone torsion angles and inter-residue geometric orientations. This term helps maintain physically realistic local geometry during optimization, preventing excessive distortion introduced by distance-driven updates. The last one is a modified FAPE loss, which measures atom-position errors after expressing coordinates in residue-local rigid frames (defined from backbone atoms, e.g., N–Cα–C). In AF-Multimer, FAPE loss is computed separately for intra-chain and inter-chain, but in the Distance-AFM, we apply a single, modified FAPE loss that operates on user-defined domains. The FAPE-loss is only computed within user-defined domains. Specifically, when computing the pairwise FAPE terms, we apply a binary mask *m_ij_* such that *m_ij_* =1 if residues *i* and *j* belong to the same domain, and *m_ij_* = 0 otherwise. As a result, all inter-domain FAPE contributions are masked out and set to zero. This intra-domain FAPE preserves local geometry and prevents the structure module from distorting well-formed intra-domain structures during iterative updates, while allowing the inter-domain (inter-chain) orientation to be primarily optimized by the distance loss *L_dis_*. The result is a multimer structure prediction that satisfies the specified distance constraints while maintaining proper inter-chain interfaces and overall complex geometry. The overall Loss function *L* is stated in Eq. 2. While a domain corresponds to an entire single chain in all our benchmark cases, users have the flexibility to define multiple distinct domains within a single chain.

Using the combined loss functions, the docking conformation undergoes iterative optimization through structure module weight updates shown in Eq. 2:

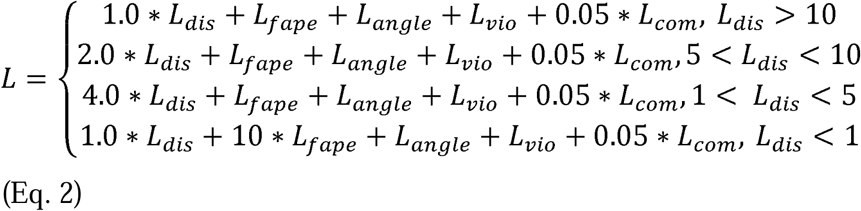

Eq. 2 explains a piecewise schedule to modulate the weight of the distance-constraint loss *L_dis_* according to how well the current structure satisfies the constraints following our previous work (Zhang et al., 2025a). When the distance loss is large, *L_dis_* >10, we keep a small weight (1.0) for *L_dis_* to avoid large updates that could severely distort intra-chain geometry. As *L_dis_* decreases to 5<*L_dis_* 10 and 1<*L_dis_* 5, we progressively increase its weight (2.0 then 4.0) so that optimization focuses on correcting inter-chain relative placement to satisfy the constraints. Once the distance constraints are almost satisfied, *L_dis_* <1, we reduce the weight back to 1.0 and upweight the structural fidelity term (10 * *L_fape_*) to restore local geometry while maintaining the already-satisfied constraints. The remaining terms (*L_angle_* and *L_vio_*) are kept unchanged across regimes to enforce stereochemistry throughout.

The development of Distance-AFM builds upon OpenFold-Multimer (Ahdritz et al., 2024) while introducing a critical preprocessing phase to address multimer-specific challenges. Details of the preprocessing algorithm, computation time, and memory usage are provided in Supplementary Information 1.

### Docking model improvement by Distance-AFM

To evaluate how well Distance-AFM can correct docking poses from incorrectly modeled complexes, we selected 27 protein complex models that were mispredicted by AFM despite high confidence scores. These protein complexes were selected from the PDB biounits, which do not contain DNA or RNA and are not too large as to exceed the GPU resources available to us. Moreover, for these complexes, AlphaFold-Multimer v2.0 failed to generate accurate models. For 21 out of 27 complexes, the DockQ docking quality score (Mirabello and Wallner, 2024) was below 0.15, and the remaining 6 targets had scores between 0.15 and 0.4. DockQ is a continuous measure of docking quality ranging from 0 to 1, where 1 represents perfect agreement with the reference structure. Models with DockQ < 0.23 are considered as Incorrect and those in the range 0.23–0.49 are of Acceptable quality. We used a cutoff of 0.15 because models with scores below this threshold typically exhibit severely incorrect inter-chain orientations rather than merely inaccurate interfaces. We therefore consider them the most challenging failure cases for AlphaFold-Multimer to recover from.

For these targets, we ran AlphaFold-Multimer v2.0 five times and selected the model with the highest ranking confidence score, 0.8 * ipTM + 0.2 * pTM, the default model-ranking metric used in AlphaFold-Multimer. ipTM (interface predicted TM-score) and pTM (predicted TM-score) are confidence estimates for the inter-chain interfaces and overall global fold, respectively. We used the top-ranked model as the starting model for refinement using Distance-AFM. This set includes 11 two-chain, 8 three-chain, and 8 four-chain complexes. Up to 12 distance constraints were used per interacting chain pair; fewer were used when fewer residue pairs were successfully matched during constraint generation. Each two-chain target used 12 constraints, whereas counts varied across chain pairs in the three- and four-chain targets. All constrained residue pairs and target distances are listed in Supplementary Table 1.

In **Figure 2** we compared Distance-AFM (D-AFM) with four representative deep-learning multimer prediction methods: AFM (Evans et al., 2022), AlphaLink2 (AL2) (Stahl et al., 2024), Chai-1 (Boitreaud et al., 2024), and Protenix (Chen et al., 2025). AFM serves as the unconstrained baseline upon which our method builds. AlphaLink2 extends AlphaFold-Multimer by incorporating experimental distance restraints (originally derived from crosslinking mass spectrometry) through a modified pair representation. Chai-1 and Protenix are recent AlphaFold3-style diffusion models, which accept user-specified contact (residue–residue distance) and pocket restraints as input features. Together, these methods provide a representative benchmark for constraint-guided complex prediction. For targets with multiple interfaces, scores were calculated for each interface and then averaged across interfaces. The data for Figure 2a–e are provided in Supplementary Table 2.

**Fig. 2.**
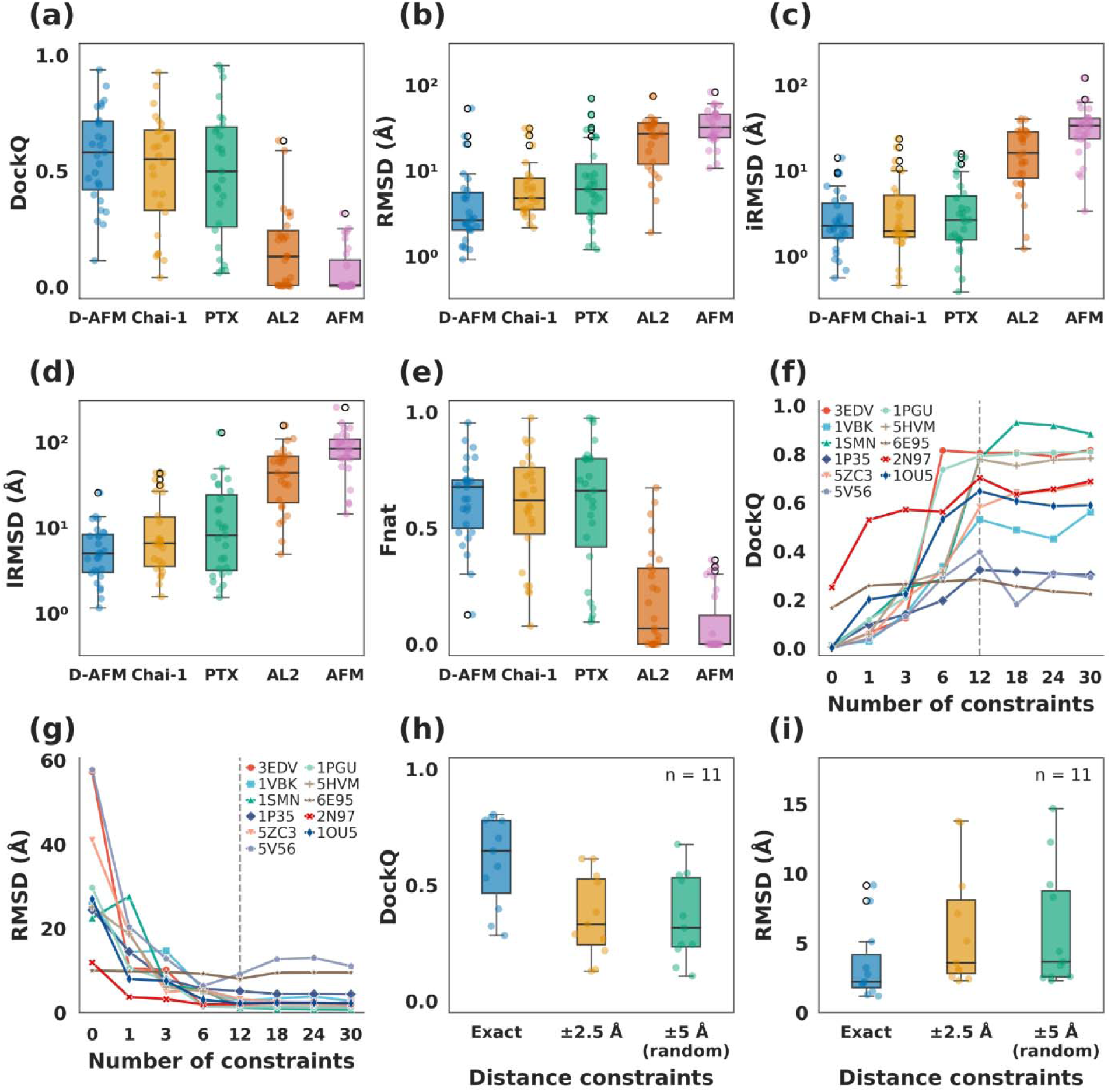
Modeling performance of Distance-AFM. A benchmark set of 27 complex targets was used. **(a–e)** Box plots showing the distribution of structural quality metrics across 27 challenging multimer targets for which AFM produces confident but incorrect predictions. The compared methods are Distance-AFM, Chai-1, Protenix, AlphaLink2 (AL2), and AFM. **(a)** DockQ scores, where values above 0.23 indicate acceptable predictions and values above 0.8 indicate high-quality predictions. The median value of each method is: D-AFM, 0.58; Chai-1, 0.55; Protenix: 0.50; AL2: 0.13; AFM: 0.01. **(b)** RMSD in Å between predicted and native structures after structural alignment. The median values of each method are: D-AFM, 2.6 Å; Chai-1, 4.8 Å; Protenix, 6.1 Å; AL2, 26.9 Å; AFM, 32.1 Å. **(c)** Interface RMSD (iRMSD), measuring the positional accuracy of interface residues. The median values of each method are: D-AFM, 2.3 Å; Chai-1, 2.0 Å; Protenix, 2.7 Å; AL2, 16.1 Å; AFM, 33.6 Å. **(d)** Ligand RMSD (lRMSD), quantifying the relative positioning and orientation of protein chains. The median values of each method are: D-AFM, 5.0 Å; Chai-1, 6.6 Å; Protenix, 8.2 Å; AL2, 43.8 Å; AFM, 83.6 Å. (e) Fraction of native contacts (Fnat), representing the proportion of inter-chain contacts preserved between predicted and native structures. The median values of each method are: D-AFM, 0.68; Chai-1, 0.62; Protenix, 0.66; AL2, 0.07; AFM, 0.00. For complexes with more than two chains, iRMSD, lRMSD, and Fnat were computed for each interface and averaged. In each box plot, the center line indicates the median, the lower and upper edges of the box indicate the first and third quartiles, respectively, and the whiskers extend to values within 1.5× the interquartile range. Points outside the whiskers are shown as outliers. All distance constraints used by Distance-AFM were derived from native structures. **(f), (g)** Model quality shown with a different number of Cα–Cα distance constraints for 11 2-chain targets (colored by PDB ID), selected from our 27-target benchmark dataset. The numbers of constraints used were 0, 1, 3, 6, 12 (our default setting; indicated by the dashed line), 18, 24, and 30. **(h)** DockQ scores for the same 11 two-chain targets using 12 Cα–Cα distance constraints under three settings: Exact, ±2.5 Å and ±5 Å (random). The median DockQ scores are 0.65, 0.33 and 0.32, respectively. **(i)** Corresponding RMSD values for the same targets and constraint settings. The median RMSDs are 2.2, 3.6 and 3.7 Å, respectively.

For evaluation, we used DockQ, the root mean square deviation (RMSD), and three component scores of DockQ, interface RMSD (iRMSD), ligand RMSD (lRMSD), and the fraction of native contacts (Fnat). Overall, D-AFM consistently demonstrates superior or near-superior performance across all metrics. In the DockQ evaluation (**Fig. 2a**), D-AFM achieves the highest median score of 0.58 among the tested methods, with a relatively tight interquartile distribution, indicating both high accuracy and stable performance across targets. Chai-1 and Protenix show intermediate performance with broader variability, whereas AL2 performs better than the baseline (AFM) but exhibits substantially lower DockQ scores than the other three methods.

The RMSD-based metrics further highlight the strength of D-AFM. D-AFM yields the lowest median RMSD (**Fig. 2b**) and lRMSD (**Fig. 2d**), and the highest median Fnat (**Fig. 2e**). Consistent with DockQ evaluation (**Fig. 2a**), AL2 displays markedly larger RMSD values, frequently extending into tens of angstroms, reflecting substantial structural deviations from native complexes. In terms of iRMSD (**Fig. 2c**), D-AFM was the second best. Overall, these results demonstrate that D-AFM provides more accurate and reliable docking predictions than the competing approaches across a broad range of evaluation criteria.

Table 1 summarizes the head-to-head comparison of D-AFM against each method. Against AF-Multimer, D-AFM wins on every target across every metric with a single exception (iRMSD on 6E95). It also dominates AlphaLink2, winning more than 90% of the 135 pairwise comparisons. Compared with Protenix, D-AFM achieved more wins in four metrics, with the exception of Fnat. The comparison with Chai-1 was the closest overall but still favored D-AFM: it achieved more wins in DockQ, RMSD, iRMSD and lRMSD, but had more losses in Fnat.

**Table 1.** Comparison of Distance-AFM and existing methods.

|  | DockQ | RMSD | iRMSD | lRMSD | Fnat |
| --- | --- | --- | --- | --- | --- |
| **Chai-1<sup>a</sup>** | 13 / 1 / 12 | 22 / 0 / 4 | 13 / 1 / 12 | 17 / 0 / 9 | 11 / 3 / 12 |
| **Protenix** | 17 / 0 / 10 | 19 / 1 / 7 | 16 / 0 / 11 | 18 / 0 / 9 | 13 / 0 / 14 |
| **Alphalink2** | 24 / 1 / 2 | 26 / 0 / 1 | 24 / 0 / 3 | 26 / 0 / 1 | 25 / 0 / 2 |
| **AF-Multimer** | 27 / 0 / 0 | 27 / 0 / 0 | 26 / 0 / 1 | 27 / 0 / 0 | 27 / 0 / 0 |

Each cell reports Win / Tie / Loss counts from the perspective of Distance-AFM against each method. A tie is defined as $|\Delta| \leq 0.01$ (DockQ, Fnat) or $|\Delta| \leq 0.01$ Å (RMSD, iRMSD, lRMSD). DockQ and Fnat are higher-is-better metrics, whereas RMSD, iRMSD, and lRMSD are lower-is-better metrics.
<sup>a</sup> Chai-1 failed to produce a prediction on target 6HCP, which is therefore excluded from the Chai-1 row ( $n = 26$ ); all other rows use $n = 27$ . Target 6HCP (2118 residues) exceeds Chai-1's hard-coded 2048-token input limit.

### Impact of Distance Constraint Density

To assess the robustness of Distance-AFM under sparse-constraint conditions, we performed an ablation study on 11 two-chain targets selected from our 27-target benchmark dataset, varying the number of imposed Cα–Cα distance constraints from 0 to 30 (Fig. 2f,g; Supplementary Table 3). The zero-constraint setting represents the unconstrained AFM baseline. For each target, constraints were randomly sampled from Cα–Cα distance pairs considering that they remain spatially well-distributed along the protein, and constraint sets of larger size were constructed as supersets of the smaller ones, so that the comparison isolates the effect of constraint density alone.

In general, DockQ increased (Fig. 2f) and RMSD decreased (Fig. 2g) as more constraints were added. The largest gains, however, were concentrated between 1 and 6 constraints, with clearly diminishing returns beyond 6. Increasing the count from 6 to 12 yielded only modest further improvement on most targets. With 6 constraints, the majority of predictions already reached the medium-to-high quality range, indicating that Distance-AFM can produce reliable structures with relatively sparse constraint information. The trend was not strictly monotonic, however. For example, for 1SMN, the RMSD increased from 22.5 Å for the AFM prediction to 27.6 Å when only a single distance restraint was applied. Although the restraint improved the interface by recovering approximately one-third of the native contacts, the overall backbone accuracy decreased slightly. A single restraint provides insufficient information to determine the correct relative orientation of the two chains, causing the model to satisfy the distance locally while adopting a distorted conformation. Using three or more restraints sufficiently constrains the relative orientation, leading to a marked improvement in accuracy.

To evaluate D-AFM’s ability to accommodate uncertainty in experimental distance measurements, we performed an ablation study using distance constraints specified as ranges rather than exact values. We extended D-AFM’s input format to accept lower and upper distance bounds and evaluated this approach using 12 constraints derived from native structures for each of 11 two-chain targets (**Fig. 2h,i**). Under the ±2.5 Å and ±5 Å (random) settings, median DockQ scores were 0.33 and 0.32, respectively, compared with 0.65 for exact constraints. Median RMSDs increased from 2.2 Å to 3.6 and 3.7 Å, respectively. Nevertheless, both interval settings produced models of at least acceptable quality (DockQ ≥ 0.23) for 8 of the 11 targets. These results demonstrate that D-AFM can accommodate distance ranges while retaining useful prediction accuracy, although performance decreases when precise distance information is unavailable. The underlying data are provided in Supplementary Table 4.

### Case studies of predicted structures

We present eight representative examples in **Figure 3**. Each panel uses four colors. The reference chain is shown in grey. The mobile chain is shown in three forms: the native structure in marine blue, the AFM prediction, the starting conformation for D-AFM is in red, and D-AFM prediction in green. We report DockQ and RMSD for both predictions. The examples are ordered by the number of chains, **Fig. 3 a–d**, two chains examples; **Fig. 3 e–g**, three chains examples; **Fig. 3 h**, four chains example.

**Fig. 3.**
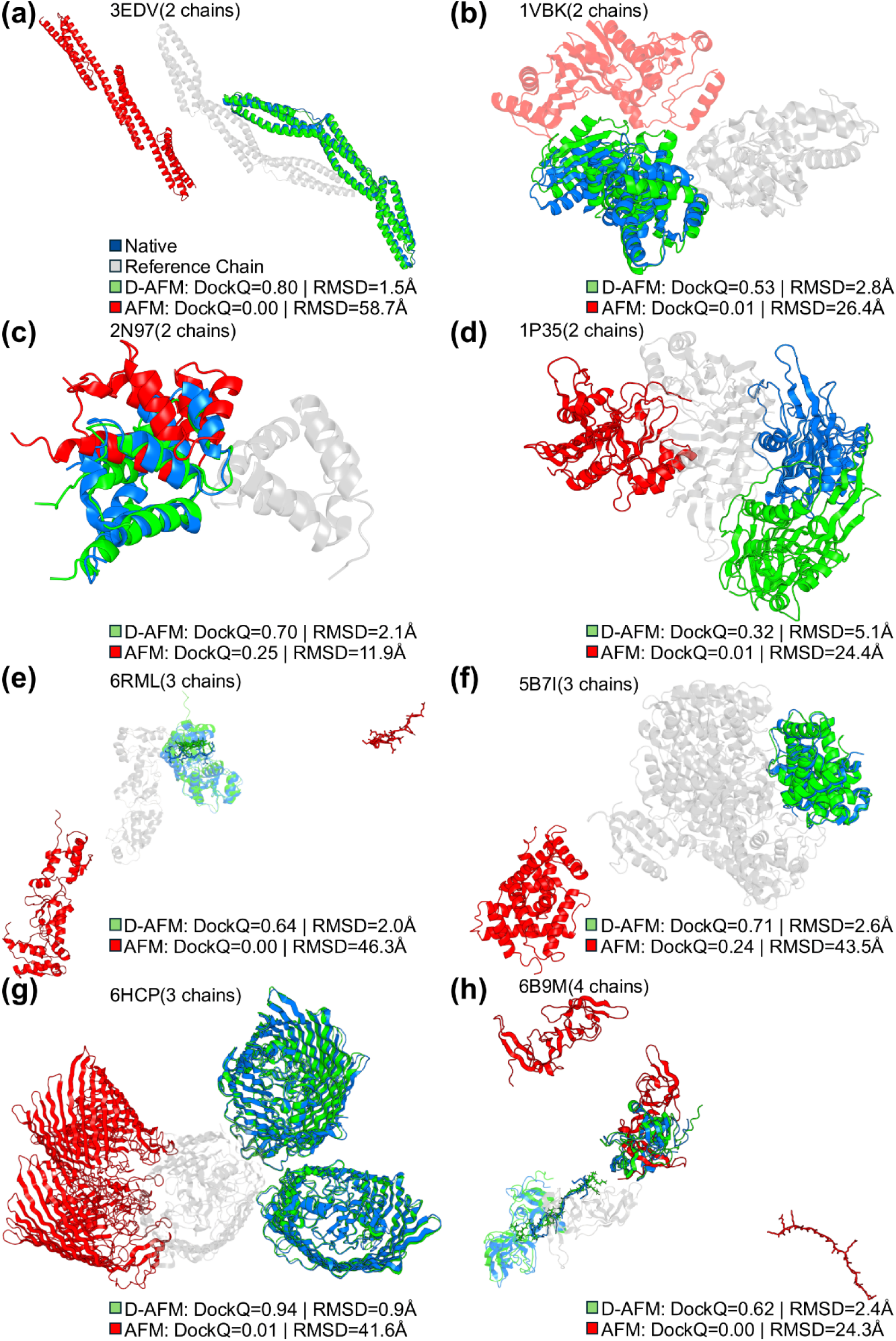
Representative complexes modeled by AlphaFold-Multimer (AFM) and Distance-AF-Multimer (D-AFM). Detailed evaluation results are available in Supplementary Table 2. Eight targets are shown, ordered by chain count (**(a)–(d)**, two chains; **(e)–(g)**, three chains; **(h)**, four chains). For each target, all structures are superimposed on a shared reference chain (grey), and the modeled position of the remaining target chain is compared against the experimental structure. Four entities are displayed: the native target chain (marine blue), the reference chain used for superposition (grey), the D-AFM model (green), and the AFM model (red). For each method, we report the DockQ score and the RMSD (Å) between the predicted and native structures after structural alignment. **(a)** Ankyrin-binding domain (repeats 14–16) of human β2-spectrin (PDB: 3EDV, 2 chains, 646 residues); D-AFM DockQ = 0.80, RMSD = 1.5 Å; AFM DockQ = 0.00, RMSD = 58.7 Å. **(b)** Putative thiamine-biosynthesis protein PH1313 from Pyrococcus horikoshii OT3 (PDB: 1VBK, 2 chains, 614 residues); D-AFM DockQ = 0.53, RMSD = 2.8 Å; AFM DockQ = 0.01, RMSD = 26.4 Å. **(c)** Death-domain (DD) homodimer of the p75 neurotrophin receptor (PDB: 2N97, 2 chains, 196 residues); D-AFM DockQ = 0.70, RMSD = 2.1 Å; AFM DockQ = 0.25, RMSD = 11.9 Å. **(d)** Baculovirus P35 caspase inhibitor from Autographa californica nucleopolyhedrovirus (PDB: 1P35, 2 chains, 598 residues); D-AFM DockQ = 0.32, RMSD = 5.1 Å; AFM DockQ = 0.01, RMSD = 24.4 Å. **(e)** TOPBP1 BRCT0,1,2 in complex with a 53BP1 phosphopeptide (PDB: 6RML, 3 chains, 599 residues); D-AFM DockQ = 0.64, RMSD = 2.0 Å; AFM DockQ = 0.00, RMSD = 46.3 Å. **(f)** Anti-CRISPR protein AcrF3 in complex with Pseudomonas aeruginosa Cas3 (PDB: 5B7I, 3 chains, 1388 residues); D-AFM DockQ = 0.71, RMSD = 2.6 Å; AFM DockQ = 0.24, RMSD = 43.5 Å. **(g)** BauA, the ferric preacinetobactin outer-membrane receptor from Acinetobacter baumannii (PDB: 6HCP, 3 chains, 2118 residues); D-AFM DockQ = 0.94, RMSD = 0.9 Å; AFM DockQ = 0.01, RMSD = 41.6 Å. **(h)** UHRF1 tandem Tudor domain (TTD) in complex with its polybasic region (PDB: 6B9M, 4 chains, 500 residues); D-AFM DockQ = 0.62, RMSD = 2.4 Å; AFM DockQ = 0.00, RMSD = 24.3 Å.

The first four panels show two-chain complexes. In the AFM prediction of 3EDV (**Fig. 3a**), the second chain is misoriented and lies far from the native interface (DockQ=0.00, RMSD=58.7 Å). D-AFM restores the correct orientation of this elongated coiled-coil and reaches DockQ=0.80 (RMSD=1.5 Å). The ablation shows that six constraints are already enough for this target. In 1VBK (**Fig. 3b**), the predicted chain has two errors at the same time. It sits at the wrong binding site, and its domain is flipped (DockQ=0.01, RMSD=26.4 Å). D-AFM corrects both and recovers a near-native interface (DockQ=0.53, RMSD=2.8 Å). 2N97 (**Fig. 3c**) shows the opposite situation. The AFM prediction has no large rigid-body error, but the interface region is positioned inaccurately, possibly because of a register shift (DockQ=0.25). This target is highly sensitive to constraints. A single constraint raises DockQ from 0.25 to 0.53, and twelve constraints reach the plateau at 0.70 (RMSD=2.1 Å). 1P35 (**Fig. 3d**) is an interesting case. When given the same constraints, other co-folding methods (Chai-1/Protenix/Alphalink2) satisfy the distances by swapping basins inside a chain instead of re-orienting the two chains. This leaves a fold-level error with RMSD around 30 Å. D-AFM keeps the internal fold of each chain and recovers an acceptable inter-chain orientation. Its DockQ is modest (0.32, RMSD=5.1 Å), but the prediction is qualitatively correct, while the competing methods are qualitatively wrong.

The next three panels show three-chain complexes. In 6RML (**Fig. 3e**), the AFM prediction has two displaced components: one full chain and a peptide are both placed away from their native positions (DockQ=0.00, RMSD=46.3 Å). D-AFM brings both back into the complex (DockQ=0.64, RMSD=2.0 Å). 5B7I (**Fig. 3f**) requires a more selective correction. This target is composed of a large receptor and a small dimer. AFM predicts the dimer correctly, but the dimer is mispositioned relative to the large receptor chain (DockQ=0.24, RMSD=43.5 Å). D-AFM moves the dimer back to its native position without disturbing its internal interface (DockQ=0.71, RMSD=2.6 Å). For 6HCP (**Fig. 3g**), the trimer prediction by AFM combines a wrong binding arrangement with a large orientation error (DockQ=0.01, RMSD=41.6 Å). D-AFM recovers an almost native structure (DockQ=0.94, RMSD=0.9 Å).

The last panel shows a complicated four-chain complex. In 6B9M (**Fig. 3h**), the AFM prediction contains several errors at once: a displaced peptide, a displaced large chain, and a rotation error (DockQ=0.00, RMSD=24.3 Å). D-AFM corrects all of them and returns the complex to a near-native arrangement (DockQ=0.62, RMSD=2.4 Å). This is the most demanding case in the set, because the method must resolve several independent errors together.

Across all eight examples, the errors range from a single misoriented chain to combined errors across four chains. This indicates that the improvement by D-AFM seen in the benchmark statistics reflects meaningful structural corrections across diverse failure types, rather than gains on a single kind of error.

### Model Selection Using Consensus-Based Scoring

Selecting the best model from candidate structures generated by Distance-AFM and other deep learning methods represents a critical challenge in the protein complex prediction. We implemented a consensus-based scoring approach of VoroIF-jury in the FTDMP framework (Olechnovic et al., 2026), which combines 20 ranking criteria derived from four scoring functions, and incorporated three additional scoring functions used in LZerD ranksum scoring (Christoffer et al., 2021). The three scores in the LZerD ranksum are knowledge-based scoring functions, GOAP (Zhou and Skolnick, 2011), DFIRE (Zhou and Zhou, 2002), and ITScorePro (Huang and Zou, 2011), which evaluate atomic interactions using statistical potentials derived from experimentally determined protein structures. Thus, the modified VoroIF-jury considers consensus of 23 ranking criteria. We used this modified VoroIF-jury because it performed very well in CASP16 complex modeling category, where our team was ranked the top 1 (Kagaya et al., 2026; Zhang et al., 2026) in the Model-1 evaluation of protein multimer prediction. The jury score ranges from 0 to 1.0, with higher values indicating stronger consensus support for a candidate model across the component rankings and structurally similar interfaces with 1.0 indicating that docking interface of the top choices of all the scores are identical.

We evaluated the scoring scheme on the same 27 targets used in **Figure 2**. The number of available models varied with complex size: the 11 two-chain targets had 4 models each, the 8 three-chain targets had 4 to 8 models each, and the 8 four-chain targets had 4 to 6 models each. For each target, we ranked the available models by their jury scores and evaluated whether the highest-ranked model corresponded to the most accurate structure. Results are summarized in **Figure 4**. Model-level results are provided in Supplementary Table 5, and component rankings and detailed Jury outputs are provided in Supplementary Table 6.

**Fig. 4.**
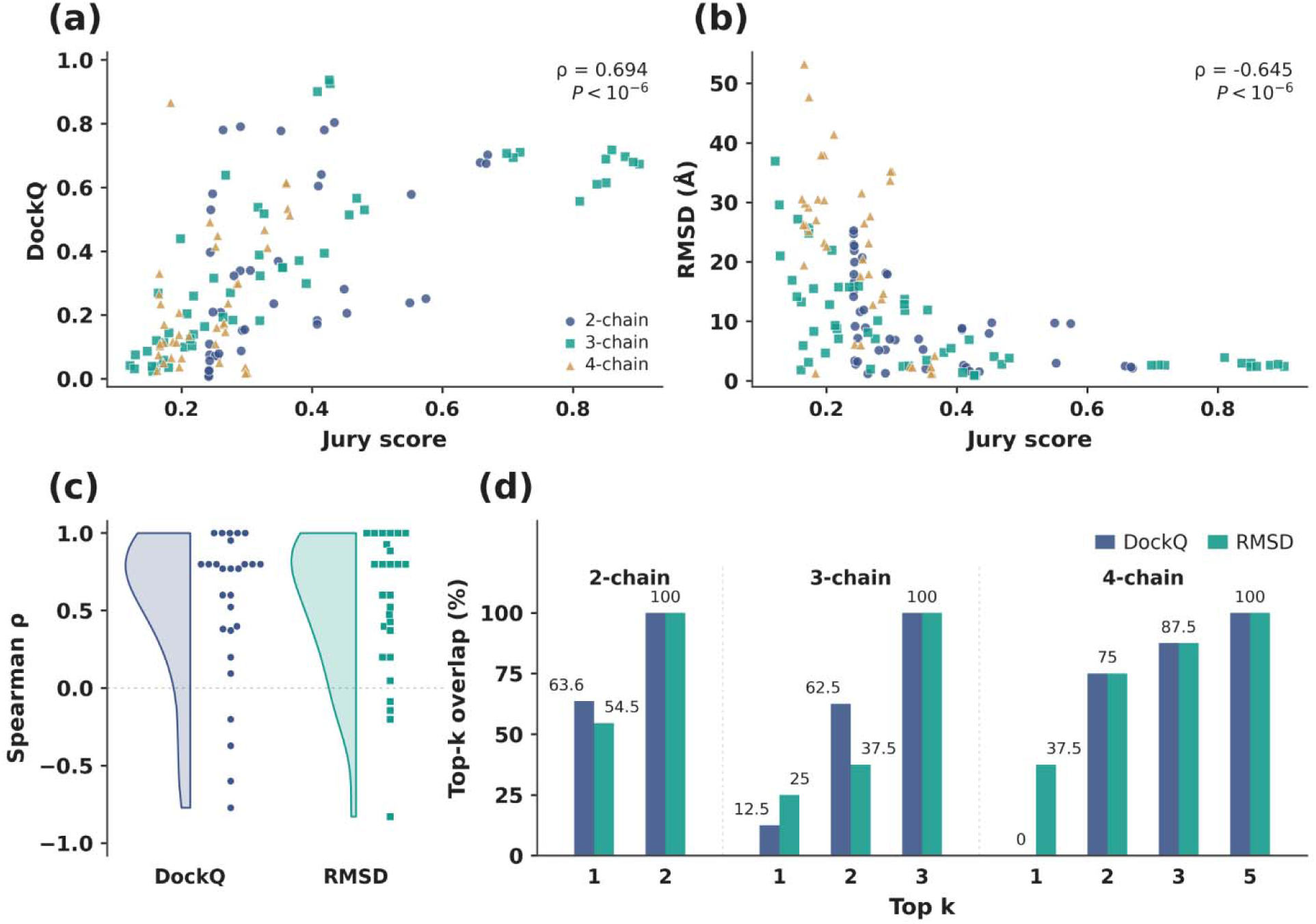
Performance evaluation of consensus-based model scoring. Consensus-based Jury scores were evaluated against native-reference DockQ and root-mean-square deviation (RMSD) for 138 candidate models across 27 protein-complex targets. Spearman’s rank correlation coefficient (ρ, ranging from −1 to 1) measures the strength and direction of the association between model rankings. P denotes the two-sided P value for the null hypothesis of zero rank correlation. **(a,b)** Jury scores versus **(a)** DockQ (ρ = 0.694, P = 4.16 × 10□²¹) and **(b)** RMSD in Å (ρ = −0.645, P = 1.35 × 10□¹□). Each scatterplot includes all 138 candidate models, with correlations calculated using global ranks. The negative correlation with RMSD indicates that higher Jury scores are associated with lower structural deviation. Navy circles, teal squares, and ochre triangles represent two-, three-, and four-chain targets, respectively. **(c)** Distributions of within-target Spearman correlations, shown as half violins with one point per target. Models were ranked by decreasing Jury score and DockQ and increasing RMSD, such that positive correlations indicate agreement between model rankings. Median ρ values were 0.771 for DockQ and 0.600 for RMSD, with positive correlations observed for 23 of the 27 targets for each metric. **(d)** Percentage of targets for which the top-k models ranked by Jury score and structural quality share at least one candidate, regardless of within-set order (ties included). For two-chain targets, Top-1 overlap was 63.6% (7/11) for DockQ and 54.5% (6/11) for RMSD, increasing to 100% (11/11) for both metrics at Top-2. For three-chain targets, Top-1 overlap was 12.5% (1/8) for DockQ and 25.0% (2/8) for RMSD; Top-2 overlap was 62.5% (5/8) and 37.5% (3/8), respectively; and Top-3 overlap reached 100% (8/8) for both metrics. For four-chain targets, Top-1 overlap was 0% (0/8) for DockQ and 37.5% (3/8) for RMSD. Top-2, Top-3, and Top-5 overlaps were 75.0% (6/8), 87.5% (7/8), and 100% (4/4), respectively, for both metrics. Top-5 analysis included only the four four-chain targets with at least five candidate models. Navy and teal indicate DockQ and RMSD, respectively, in (c) and (d). Source data are provided in Supplementary Tables 5 and 6.

**Fig. 4a and Fig. 4b show** the distribution of DockQ and RMSD, respectively, relative to the jury score for all 138 models from the 27 targets. Jury scores showed a strong positive correlation with DockQ (Spearman’s ρ = 0.694, P-value = 4.16 × 10□²¹) and a strong negative correlation with RMSD (ρ = −0.645, P-value = 1.35 × 10□¹□), indicating that higher Jury scores were associated with more accurate structural models. We next examined how well Jury scores ranked candidate models within individual targets (**Fig. 4c**). The median Spearman’s rank correlation coefficients were 0.771 for DockQ and 0.600 for RMSD, with positive correlations observed for 23 of the 27 targets for each metric. These results indicate that Jury scores generally reflect differences in structural accuracy among candidate models for the same target.

We next evaluated Jury scoring by measuring whether the top-k candidates ranked by Jury score and structural quality shared at least one model, regardless of within-set order (Fig. 4d). For two-chain targets, Top-1 overlap was 63.6% for DockQ and 54.5% for RMSD, increasing to 100% for both metrics at Top-2. For three-chain targets, Top-1 overlap was 12.5% for DockQ and 25.0% for RMSD, and Top-3 overlap reached 100% for both metrics. For four-chain targets, Top-1 overlap was 0% for DockQ and 37.5% for RMSD; Top-2 and Top-3 overlaps were 75.0% and 87.5%, respectively, for both metrics. Thus, overlap between Jury-ranked and quality-ranked candidate sets increased as more candidates were considered.

### Modeling peptide-receptor docking

We further modelled peptide-receptor complex targets using Distance-AFM. Peptide docking is particularly prone to placement and orientation errors because short peptides are conformationally flexible and form comparatively small interfaces (Ling et al., 2026). We selected six targets for which AFM produced incorrect peptide poses, as shown in the work with DistPepFold (Zhang et al., 2025b).

Six constraints were used for each peptide, resulting in twelve constraints for 5AER, which contains two peptides, and six constraints for each of the other five targets. Target definitions and constraints are provided in Supplementary Table 7. The selected targets covered five qualitatively distinct failure modes: a spatially detached peptide (1CN3); a register-shifted peptide (8U51); binding at an incorrect receptor site (8G4Y and 9CK0); reversal of the peptide N-to-C orientation (7ZFR); and a composite failure combining N-to-C inversion with incorrect binding-site placement in a complex containing two peptides (5AER). Because D-AFM uses DistPepFold-derived weights in the peptide setting, we expected it to be well suited to peptide correction. We therefore used this peptide–protein set as a targeted correction stress test, asking whether constraint-guided refinement could redirect one or more mispredicted peptide chains towards their native binding modes.

Across the six targets, D-AFM improved DockQ and Fnat and reduced the peptide-backbone RMSD (ligand RMSD), for every target (**Fig. 5a–f**). DockQ and Fnat were calculated for each peptide-containing interface and, when multiple peptides were present, averaged across peptides; this occurred only for 5AER. Ligand RMSD was calculated over matched peptide backbone atoms (N, Cα, C, and O) after superposition of the shared receptor backbone atoms.

**Fig. 5.**
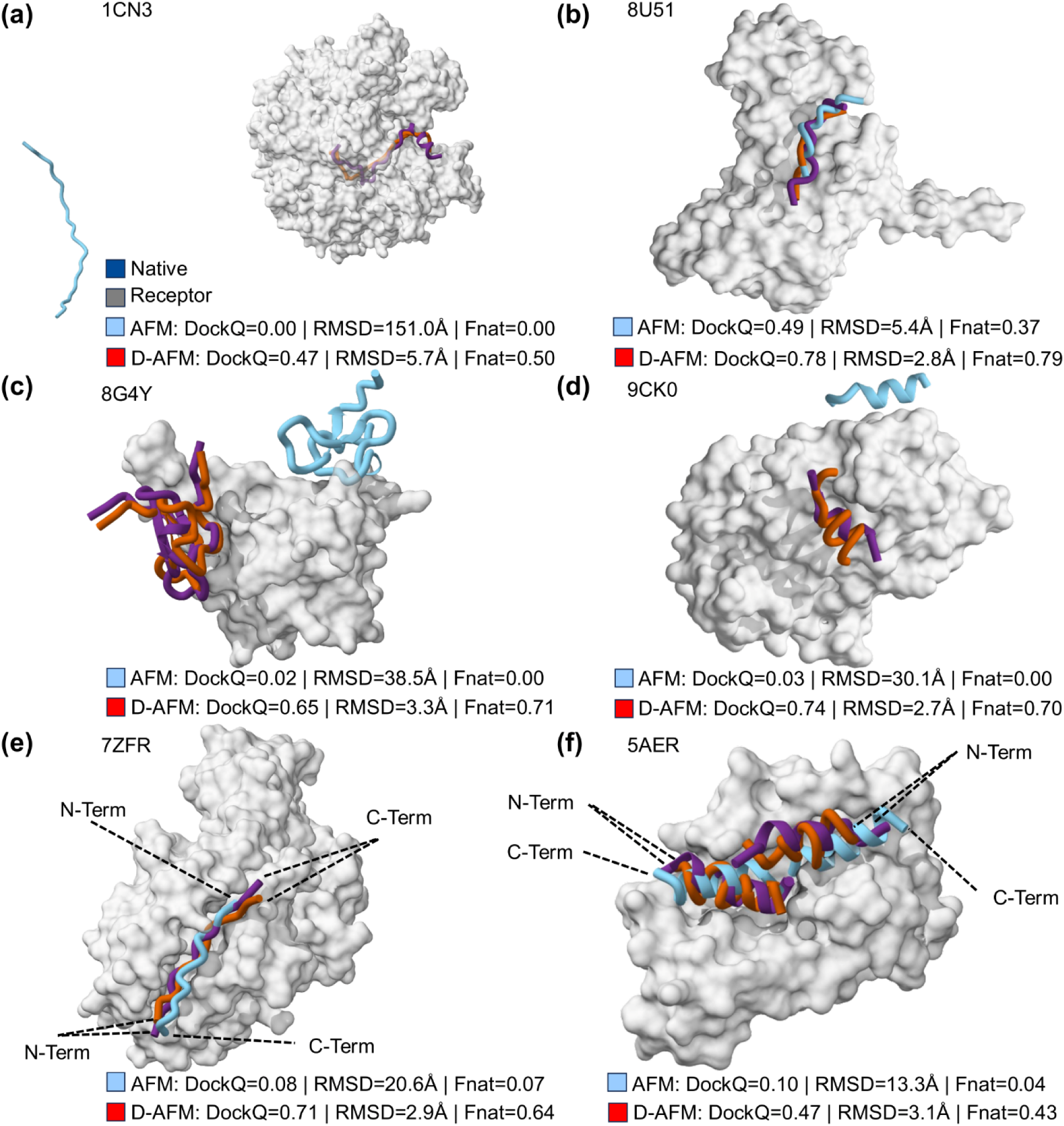
Structure correction for peptide-protein complexes. AFM and D-AFM predictions are compared for six peptide–protein complexes. For each target, the AFM and D-AFM models were superimposed on the native structure using the receptor for alignment (grey). The remaining components, including the peptide chain(s), are shown for the native structure (marine blue), AFM model (light blue), and D-AFM model (red). DockQ and Fnat are reported as mean values across all interfaces involving at least one peptide chain; receptor-only interfaces are excluded. RMSD is calculated over the matched peptide backbone atoms (N, Cα, C, and O) after receptor superposition (i.e. ligand RMSD). **(a)** Polyomavirus major capsid protein VP1 pentamer bound to a C-terminal fragment of minor capsid protein VP2 (PDB: 1CN3, 6 chains, 1,444 residues); D-AFM DockQ = 0.47, RMSD = 5.7 Å, Fnat = 0.50; AFM DockQ = 0.00, RMSD = 151.0 Å, Fnat = 0.00. **(b)** Selected Klebsiella pneumoniae family 1 encapsulin shell protomer bound to the targeting-peptide segment of a SUMO–targeting peptide fusion protein (PDB: 8U51, 2 chains, 278 residues); D-AFM DockQ = 0.78, RMSD = 2.8 Å, Fnat = 0.79; AFM DockQ = 0.49, RMSD = 5.4 Å, Fnat = 0.37. **(c)** Human E3 ubiquitin-protein ligase ZNRF3 extracellular domain bound to the synthetic peptide MK1-3.6.10 (PDB: 8G4Y, 2 chains, 217 residues); D-AFM DockQ = 0.65, RMSD = 3.3 Å, Fnat = 0.71; AFM DockQ = 0.02, RMSD = 38.5 Å, Fnat = 0.00. **(d)** Human peroxisome proliferator-activated receptor gamma (PPARγ) ligand-binding domain bound to an NF-κB inhibitor beta (IκBβ/NFKBIB)-derived peptide (PDB: 9CK0, 2 chains, 299 residues); D-AFM DockQ = 0.74, RMSD = 2.7 Å, Fnat = 0.70; AFM DockQ = 0.03, RMSD = 30.1 Å, Fnat = 0.00. **(e)** Human HLA-DP (DPA102:01–DPB101:01) αβ heterodimer in complex with a synthetic 12-residue peptide bound in the reverse direction (PDB: 7ZFR, 3 chains, 542 residues); D-AFM DockQ = 0.71, RMSD = 2.9 Å, Fnat = 0.64; AFM DockQ = 0.08, RMSD = 20.6 Å, Fnat = 0.07. **(f)** Rat neuronal calcium sensor-1 in complex with two copies of a synthetic human dopamine D2 receptor-derived peptide (PDB: 5AER, 3 chains, 218 residues); D-AFM DockQ = 0.47, RMSD = 3.1 Å, Fnat = 0.43; AFM DockQ = 0.10, RMSD = 13.3 Å, Fnat = 0.04.

We first examined large spatial displacement and rotational misorientation in AFM models. In 1CN3 (**Fig. 5a**), AFM placed the peptide far from the VP1 pentamer receptor and in an incorrect orientation (DockQ = 0.00, RMSD = 151.0 Å and Fnat = 0.00). D-AFM corrected the fragment’s rotational misorientation and repositioned it within the central axial cavity of the VP1 pentamer, recovering the characteristic insertion of the VP2 C-terminal segment into the pentameric assembly. This correction increased DockQ to 0.47 and Fnat to 0.50 while reducing RMSD to 5.7 Å.

We next examined errors confined to or near the receptor interface. In 8U51 (**Fig. 5b**), AFM positioned the peptide in approximately the correct groove but with a register shift. D-AFM increased DockQ from 0.49 to 0.78 and Fnat from 0.37 to 0.79, whereas RMSD changed only from 5.4 to 2.8 Å. This disproportionate gain in interface-sensitive metrics relative to RMSD is consistent with correction of the local register without a large rigid-body movement. For the next target (**Fig. 5c**), 8G4Y, AFM placed the peptide at an incorrect receptor region, yielding DockQ = 0.02, RMSD = 38.5 Å and Fnat = 0.00. D-AFM redirected the peptide towards the native binding region, increasing DockQ to 0.65 and Fnat to 0.71 while reducing RMSD to 3.3 Å. Similarly, in 9CK0 (**Fig. 5d**), AFM placed the peptide at an incorrect binding site (DockQ = 0.03, RMSD = 30.1 Å and Fnat = 0.00). D-AFM relocated the peptide to the native binding region, increasing DockQ to 0.74 and Fnat to 0.70 while reducing RMSD to 2.7 Å (**Fig. 5d**). Together, these cases illustrate correction of both local register errors and larger binding-site misassignments.

The two final cases introduced directionality errors, either alone or coupled to incorrect placement. In 7ZFR (**Fig. 5e**), AFM reversed the peptide N-to-C orientation. D-AFM moved the peptide towards the native orientation, increasing DockQ from 0.08 to 0.71 and Fnat from 0.07 to 0.64 while reducing RMSD from 20.6 to 2.9 Å. The 5AER complex, which contains two peptides, represented the most complex failure mode in this set, combining N-to-C inversion with incorrect binding-site placement. D-AFM increased DockQ from 0.10 to 0.47 and Fnat from 0.04 to 0.43 and reduced RMSD from 13.3 to 3.1 Å (**Fig. 5f**). These directionality gains suggest that the full D-AFM pipeline can exploit orientational information beyond approximate pocket location. Nevertheless, the final DockQ and Fnat values for 5AER indicate that correction remained incomplete in this multi-peptide complex.

Together, these six selected cases indicate that D-AFM correction was not limited to translating detached peptides; it improved predictions involving receptor orientation, peptide register, binding-site selection and N-to-C directionality, including a complex containing two peptides. The DistPepFold-derived weights may contribute a useful peptide-specific prior, but the present comparison does not isolate their effect from that of the distance constraints or other components of D-AFM. Moreover, because the targets were deliberately selected for peptide-pose failures, these results illustrate the range of error types that can be improved rather than providing an unbiased estimate of D-AFM performance across peptide–protein complexes.

### Modeling large protein complexes

Finally, we show modeling results for five large complex structures containing five to ten chains (Fig. 6). Target definitions and constraints are provided in Supplementary Table 8. For each example, we show an intermediate structure from Phase 2 and the corresponding final structure after Phase 3 refinement.

**Fig. 6.**
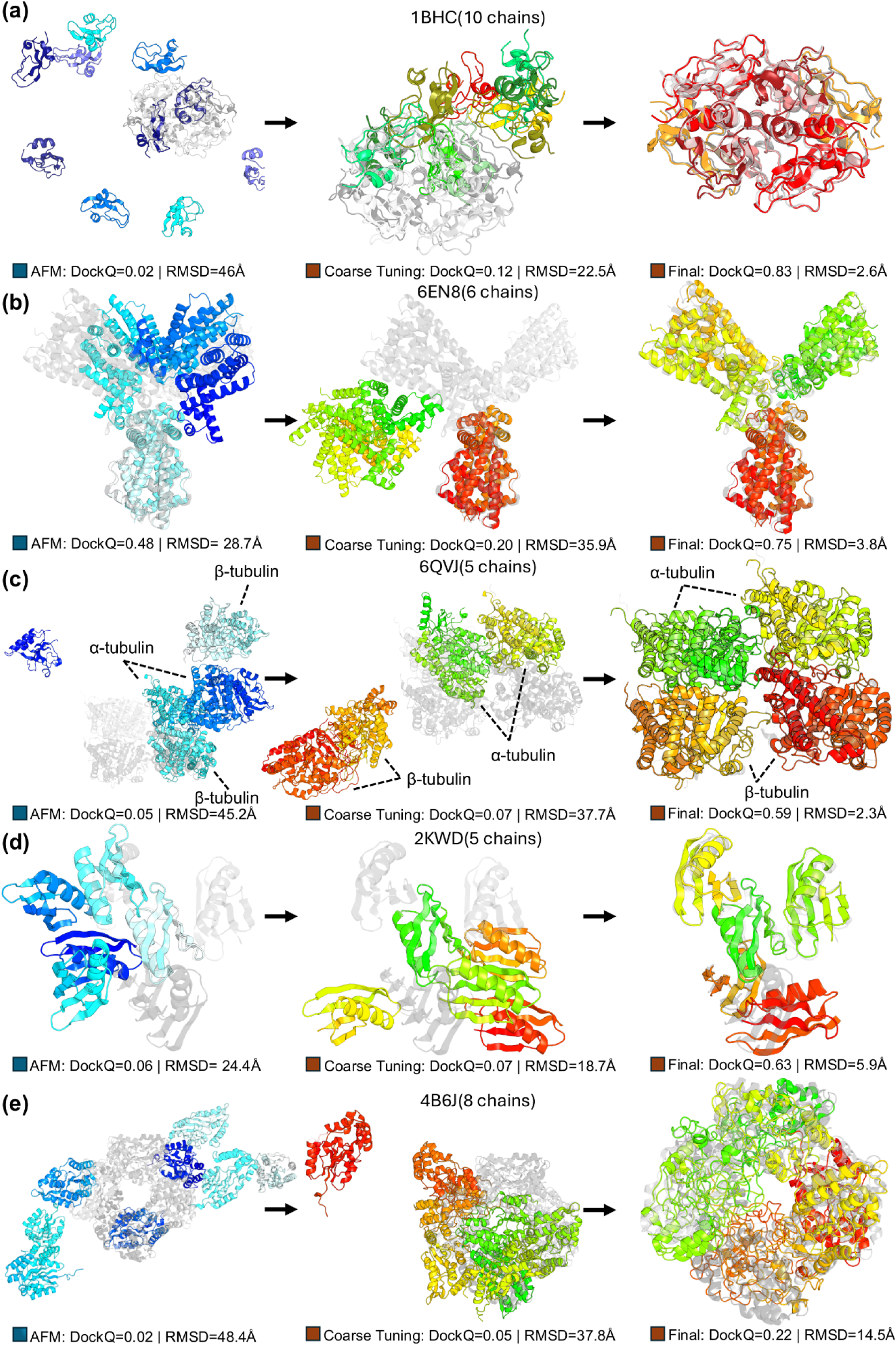
Examples of large multimeric protein assemblies. Five representative large multimeric assembly targets are shown, with each row displaying the initial AFM model (left), the intermediate structure after coarse tuning (middle), and the final D-AFM model (right). Experimental structures are shown in gray, initial AFM models in shades of blue and cyan, and intermediate and final models in shades of red to green. Arrows indicate progression through the two modeling phases. **(a)** Bovine pancreatic trypsin inhibitor (PDB: 1BHC, 10 chains, 580 residues); AFM DockQ = 0.02, RMSD = 46.0 Å; coarse tuning: DockQ = 0.12, RMSD = 22.5 Å; final structure DockQ = 0.83, RMSD = 2.6 Å. **(b)** TetR-family transcriptional regulator SaFadR from Sulfolobus acidocaldarius (PDB: 6EN8, 6 chains, 1176 residues); AFM DockQ = 0.48, RMSD = 28.7 Å; coarse tuning DockQ = 0.20, RMSD = 35.9 Å; final structure DockQ = 0.75, RMSD = 3.8 Å. **(c)** Human CAMSAP1 CKK domain in complex with α- and β- tubulin (PDB: 6QVJ, 5 chains, 1964 residues); AFM DockQ = 0.05, RMSD = 45.2 Å; coarse tuning DockQ = 0.07, RMSD = 37.7 Å; final structure DockQ = 0.59, RMSD = 2.3 Å. **(d)** B1 domain of immunoglobulin G-binding protein G (GB1) (PDB: 2KWD, 5 chains, 280 residues); AFM DockQ = 0.06, RMSD = 24.4 Å; coarse tuning DockQ = 0.07, RMSD = 18.7 Å; final structure DockQ = 0.63, RMSD = 5.9 Å. **(e)** Phosphoserine phosphatase from Thermococcus onnurineus (PDB: 4B6J, 8 chains, 1608 residues); AFM DockQ = 0.02, RMSD = 48.4 Å; coarse tuning DockQ = 0.05, RMSD = 37.8 Å; final structure DockQ = 0.22, RMSD = 14.5 Å.

The first example is bovine pancreatic trypsin inhibitor (PDB ID: 1BHC), a decameric complex containing 580 amino acids (**Fig. 6a**). In the initial AFM model, only two chains out of ten were in contact, while the remaining chains were dispersed. We generated five coarse-tuned models, each of which was refined in Phase 3. The intermediate model brought all ten chains into contact, although their relative orientations remained inaccurate and the RMSD was still 22.5 Å. Fine tuning then improved chain orientations and interface packing, producing a near-native assembly with a DockQ score of 0.83 and an RMSD of 2.6 Å. In this case, coarse tuning established the overall assembly, and fine tuning corrected how the chains fitted together.

The second example (**Fig. 6b**) is the SaFadR assembly (PDB ID: 6EN8), a six-chain complex comprising three dimeric units. In the initial AFM model, the three dimers were formed, but their relative orientations differed from those in the experimental structure. Coarse tuning disrupted this arrangement, and DockQ decreased from 0.48 to 0.20. The final D-AFM model subsequently recovered the overall assembly and improved the interfaces, achieving an RMSD of 3.8 Å and DockQ of 0.75 relative to the experimental structure. Thus, the temporary loss of interface accuracy did not prevent successful recovery. This trajectory suggests that coarse tuning provides an alternative starting arrangement from which fine tuning can rebuild interfaces using the structural preferences learned by AFM.

Recovery also extended to the heteromeric CAMSAP1–tubulin complex (PDB 6QVJ), comprising one CAMSAP1 CKK domain, two α-tubulin chains, and two β-tubulin chains (**Fig. 6c**). The initial AFM model contained two weakly formed α/β-tubulin interfaces, with DockQ scores of 0.23 and 0.24, but failed to assemble the complete complex. Coarse tuning disrupted these initial interfaces while rearranging the chains. Fine tuning then recovered all eight native interfaces, yielding an overall DockQ score of 0.59 and an RMSD of 2.3 Å. This example shows that recovery can involve rebuilding initially imperfect interfaces as part of a larger heteromeric assembly.

The final two examples illustrate cases in which Distance-AFM substantially improved the overall assembly, although some chains remained incorrectly oriented in the final models. The fourth example is a five-chain complex of Immunoglobulin G-binding protein G (PDB ID: 2KWD) (**Fig. 6d**). The initial AFM model showed a totally incorrect arrangement of all the five chains, with an RMSD of 24.4 Å and a DockQ score of 0.06. Coarse tuning reduced RMSD to 18.7 Å by bringing chain D closer to the correct position, although its orientation remained incorrect. Fine tuning subsequently recovered the placement of four of the five chains. The final model reached a DockQ score of 0.63 and an RMSD of 5.9 Å. However, chain E (red) remained misoriented by approximately 72°, despite being positioned correctly relative to the other chains. Nevertheless, its distance constraints showed a mean deviation of only 1.16 Å. A rigid-body rotation scan confirmed that substantially different orientations could yield similarly small distance deviations, indicating that the selected constraints provided insufficient orientational information to resolve the remaining error.

The final example is the eight-chain phosphoserine phosphatase homoctamer (PDB 4B6J; 1,608 residues), which improved substantially but remained challenging (**Fig. 6e**). AFM produced a model with an RMSD of 48.4 Å. Coarse tuning in D-AFM brought more chains together, reducing the RMSD to 37.8 Å and improving two of the 16 native interfaces toward acceptable quality, although most remained inaccurate. The final D-AFM model further reduced the RMSD to 14.5 Å and brought most chains closer to their native arrangement. However, residual orientation errors and intrachain deformation remained, with monomer Cα RMSD increasing from 3.4 Å in the AFM model to 4.7 Å in the final model. The final DockQ score was 0.22, just below the acceptable-quality threshold of 0.23.

Together, these examples show that Distance-AFM can recover multichain assemblies, including heteromeric complexes, from incorrect AFM predictions. Coarse tuning explores alternative chain arrangements that seed fine tuning, while partially recovered cases reveal limitations from ambiguous orientations and residual inter- and intrachain errors.

## Discussion

We developed Distance-AFM, an extension of Distance-AF (Zhang et al., 2025a) that incorporates inter-chain distance constraints into protein complex structure prediction. Protein structure prediction methods are now routinely used by experimental biologists to guide experimental design and interpret biological data. However, although these methods often produce highly accurate models, there is currently no straightforward way to incorporate experimental evidence when the predicted structures disagree with known observations. Recent approaches, such as Chai-1 (Boitreaud et al., 2024) and Protenix (Chen et al., 2025), address this limitation by retraining the prediction network to handle structural constraints, but doing so requires large-scale training datasets and substantial computational resources. In contrast, Distance-AFM incorporates distance constraints through iterative optimization of a pre-trained AlphaFold-Multimer model, requiring no retraining and only modest additional computation for each target. Despite this lightweight framework, Distance-AFM achieves competitive performance while enabling flexible integration of experimental restraints.

Sparse distance restraints of this kind are readily obtained from experimental techniques such as XL-MS (Lee and O’Reilly, 2023), cryo-EM, NMR, and FRET (Peter et al., 2022). Moreover, the restraints need not originate from experiments; users can also specify hypothetical contacts to evaluate the structural feasibility of proposed interaction interfaces. This flexibility makes Distance-AFM a practical tool for studying transient, flexible, or otherwise difficult-to-characterize protein complexes, where comprehensive experimental data are often unavailable.

Future work will extend Distance-AFM to integrate a broader range of experimental restraints beyond pairwise distances. We plan to incorporate volumetric information derived from SAXS (Hura et al., 2019), cryo-EM (Li et al., 2025; Terashi et al., 2024), and cryo-electron tomography (cryo-ET), as well as restraints from hydrogen–deuterium exchange mass spectrometry (HDX-MS) (Chen et al., 2023; Tran et al., 2025; Wang et al., 2025), mutagenesis, and other biophysical experiments.

Based on Distance-AFM, we also aim to develop adaptive strategies for selecting the most informative restraints when experimental data are sparse or acquisition costs are limited, while exploring integration with emerging protein structure prediction architectures without sacrificing the training-free optimization paradigm. Ultimately, we envision Distance-AFM as a flexible framework for next-generation integrative structural modeling (Hancock et al., 2022), where complementary experimental evidence and AI-based prediction are combined to solve structures beyond the reach of a single AF or AFM prediction. This capability will enable the modeling of increasingly large and complex biological assemblies, including membrane protein supercomplexes, multienzyme complexes, chromatin-associated assemblies, viral replication and capsid complexes, and other cell-scale macromolecular systems.

## Supporting information

Supplemental Information Doc

## Supplementary Material

Supplementary material is provided in two files. The supplementary document (dafm_suppl_v1.3.docx) describes the starting-conformation generation algorithm, computation time and memory usage, and the supplementary table legends. The accompanying workbook (suppl_table_v1.1.xlsx) contains Supplementary Tables 1–8, including benchmark target definitions and distance constraints, model-quality comparisons, constraint-number and uncertainty analyses, results of consensus scoring, and target definitions and constraints for peptide-protein complexes and large multimeric assemblies.

## Author Contributions

DK conceived and supervised the study, acquired funding, KL developed the method and software, and performed the benchmark study. KL and DK data analysis, KL wrote the initial manuscript. YZ provided guidance on adapting Distance-AF. ZZ provided DistPepFold model weights, AlphaFold-related scripts, and peptide–protein test cases. TN was involved in benchmark dataset selection. DK critically edited the manuscript. All authors approved the manuscript.

## Data Availability Statement

Benchmark target information, distance constraints, and evaluation results are provided in the Supplementary Material. The Distance-AF Multimer source code and model weights are available at https://github.com/kiharalab/Distance-AFM

## AI Disclosure Statement

ChatGPT (OpenAI) was used to help edit the manuscript based on the initial writing and to check the code and data. The suggestions were checked against the original text, scripts, source data, and results and reviewed by the lead author. The authors take full responsibility for the manuscript.

## References

Aderinwale T, Christoffer CW, Sarkar D, Alnabati E, Kihara D (2020) Computational structure modeling for diverse categories of macromolecular interactions. Current Opinion in Structural Biology 64:1–8.

Ahdritz G, Bouatta N, Floristean C, Kadyan S, Xia Q, Gerecke W, O’Donnell TJ, Berenberg D, Fisk I, Zanichelli N, Zhang B, Nowaczynski A, Wang B, Stepniewska-Dziubinska MM, Zhang S, Ojewole A, Guney ME, Biderman S, Watkins AM, Ra S, Lorenzo PR, Nivon L, Weitzner B, Ban YA, Chen S, Zhang M, Li C, Song SL, He Y, Sorger PK, Mostaque E, Zhang Z, Bonneau R, AlQuraishi M (2024) OpenFold: retraining AlphaFold2 yields new insights into its learning mechanisms and capacity for generalization. Nature Methods 21:1514–1524.

Boitreaud J, Dent J, McPartlon M, Meier J, Reis V, Rogozhnikov A, Wu K. Chai-1: Decoding the molecular interactions of life. (2024) bioRxiv.

Chen X, Wang Y, Xu Z, Cheng ML, Ma QQ, Li RT, Wang ZJ, Zhao H, Zuo X, Li XF, Fang X, Qin CF (2023) Zika virus RNA structure controls its unique neurotropism by bipartite binding to Musashi-1. Nature Communications 14:1134.

Chen X, Zhang Y, Lu C, Ma W, Guan J, Gong C, Yang J, Zhang H, Zhang K, Wu S, Zhou K, Yang Y, Liu Z, Wang L, Shi B, Shi S, Xiao W. Protenix - Advancing Structure Prediction Through a Comprehensive AlphaFold3 Reproduction. (2025) bioRxiv.

Christoffer C, Chen S, Bharadwaj V, Aderinwale T, Kumar V, Hormati M, Kihara D (2021) LZerD webserver for pairwise and multiple protein-protein docking. Nucleic Acids Res 49:W359–W365.

de Vries SJ, van Dijk M, Bonvin AM (2010) The HADDOCK web server for data-driven biomolecular docking. Nature Protocols 5:883–897.

Evans R, O’Neill M, Pritzel A, Antropova N, Senior A, Green T, Žídek A, Bates R, Blackwell S, Yim J, Ronneberger O, Bodenstein S, Zielinski M, Bridgland A, Potapenko A, Cowie A, Tunyasuvunakool K, Jain R, Clancy E, Kohli P, Jumper J, Hassabis D. Protein complex prediction with AlphaFold-Multimer. (2022) bioRxiv.

Hancock M, Peulen TO, Webb B, Poon B, Fraser JS, Adams P, Sali A (2022) Integration of software tools for integrative modeling of biomolecular systems. J Struct Biol 214:107841.

Harini K, Christoffer C, Gromiha MM, Kihara D (2023) Pairwise and Multi-chain Protein Docking Enhanced Using LZerD Web Server. Methods Mol Biol 2690:355–373.

Huang SY, Zou X (2011) Statistical mechanics-based method to extract atomic distance-dependent potentials from protein structures. Proteins 79:2648–2661.

Hura GL, Hodge CD, Rosenberg D, Guzenko D, Duarte JM, Monastyrskyy B, Grudinin S, Kryshtafovych A, Tainer JA, Fidelis K, Tsutakawa SE (2019) Small angle X-ray scattering-assisted protein structure prediction in CASP13 and emergence of solution structure differences. Proteins 87:1298–1314.

Jumper J, Evans R, Pritzel A, Green T, Figurnov M, Ronneberger O, Tunyasuvunakool K, Bates R, Zidek A, Potapenko A, Bridgland A, Meyer C, Kohl SAA, Ballard AJ, Cowie A, Romera-Paredes B, Nikolov S, Jain R, Adler J, Back T, Petersen S, Reiman D, Clancy E, Zielinski M, Steinegger M, Pacholska M, Berghammer T, Bodenstein S, Silver D, Vinyals O, Senior AW, Kavukcuoglu K, Kohli P, Hassabis D (2021) Highly accurate protein structure prediction with AlphaFold. Nature 596:583–589.

Kagaya Y, Nakamura T, Verburgt J, Jain A, Terashi G, Punuru P, Tugolukova E, Park JH, Saha A, Huang D, Kihara D (2026) Structure Modeling Protocols for Protein Multimer and RNA in CASP16 With Enhanced MSAs, Model Ranking, and Deep Learning. Proteins 94:167–182.

Kozakov D, Hall DR, Xia B, Porter KA, Padhorny D, Yueh C, Beglov D, Vajda S (2017) The ClusPro web server for protein-protein docking. Nature Protocols 12:255–278.

Lee K, O’Reilly FJ (2023) Cross-linking mass spectrometry for mapping protein complex topologies in situ. Essays Biochem 67:215–228.

Li S, Terashi G, Zhang Z, Kihara D (2025) Advancing structure modeling from cryo-EM maps with deep learning. Biochem Soc Trans 53:259–265.

Ling K, Li S, Zhang Z, Shin WH, Kihara D (2026) Peptide-protein docking: from physics-based models to generative intelligence. Chemical Communications 62:7235–7247.

Lu D, Yu S, Huang Y, Gong X (2025) Multimeric protein interaction and complex prediction: Structure, dynamics and function. Computational and Structural Biotechnology Journal 27:1975–1997.

Lyskov S, Gray JJ (2008) The RosettaDock server for local protein-protein docking. Nucleic Acids Res 36:W233–238.

Mirabello C, Wallner B (2024) DockQ v2: improved automatic quality measure for protein multimers, nucleic acids, and small molecules. Bioinformatics 40.

Mirdita M, von den Driesch L, Galiez C, Martin MJ, Soding J, Steinegger M (2017) Uniclust databases of clustered and deeply annotated protein sequences and alignments. Nucleic Acids Res 45:D170–D176.

Morcos F, Pagnani A, Lunt B, Bertolino A, Marks DS, Sander C, Zecchina R, Onuchic JN, Hwa T, Weigt M (2011) Direct-coupling analysis of residue coevolution captures native contacts across many protein families. Proc Natl Acad Sci U S A 108:E1293–1301.

Olechnovic K, Banciul R, Dapkunas J, Venclovas C (2026) FTDMP: A Framework for Protein-Protein, Protein-DNA, and Protein-RNA Docking and Scoring. Proteins 94:1753–1761.

Passaro S, Corso G, Wohlwend J, Reveiz M, Thaler S, Somnath VR, Getz N, Portnoi T, Roy J, Stark H, Kwabi-Addo D, Beaini D, Jaakkola T, Barzilay R. Boltz-2: Towards Accurate and Efficient Binding Affinity Prediction. (2025) bioRxiv.

Peter MF, Gebhardt C, Machtel R, Munoz GGM, Glaenzer J, Narducci A, Thomas GH, Cordes T, Hagelueken G (2022) Cross-validation of distance measurements in proteins by PELDOR/DEER and single-molecule FRET. Nature Communications 13:4396.

Richardson L, Allen B, Baldi G, Beracochea M, Bileschi ML, Burdett T, Burgin J, Caballero-Perez J, Cochrane G, Colwell LJ, Curtis T, Escobar-Zepeda A, Gurbich TA, Kale V, Korobeynikov A, Raj S, Rogers AB, Sakharova E, Sanchez S, Wilkinson DJ, Finn RD (2023) MGnify: the microbiome sequence data analysis resource in 2023. Nucleic Acids Res 51:D753–D759.

Stahl K, Warneke R, Demann L, Bremenkamp R, Hormes B, Brock O, Stulke J, Rappsilber J (2024) Modelling protein complexes with crosslinking mass spectrometry and deep learning. Nature Communications 15:7866.

Steinegger M, Soding J (2018) Clustering huge protein sequence sets in linear time. Nature Communications 9:2542.

Terashi G, Wang X, Prasad D, Nakamura T, Kihara D (2024) DeepMainmast: integrated protocol of protein structure modeling for cryo-EM with deep learning and structure prediction. Nature Methods 21:122–131.

Tran MH, Martina CE, Moretti R, Nagel M, Schey KL, Meiler J (2025) RosettaHDX: Predicting antibody-antigen interaction from hydrogen-deuterium exchange mass spectrometry data. J Struct Biol 217:108166.

UniProt C (2025) UniProt: the Universal Protein Knowledgebase in 2025. Nucleic Acids Res 53:D609–D617.

Venkatraman V, Yang YD, Sael L, Kihara D (2009) Protein-protein docking using region-based 3D Zernike descriptors. BMC bioinformatics 10:407.

Wang L, Tucs A, Ding S, Tsuda K, Sljoka A (2025) HDXRank: A Deep Learning Framework for Ranking Protein Complex Predictions with Hydrogen-Deuterium Exchange Data. J Chem Theory Comput 21:7173–7187.

Zhang J, Yuan R, Kryshtafovych A, Pei J, Kretsch RC, Schaeffer RD, Zhou J, Das R, Grishin NV, Cong Q (2026) Assessment of Protein Complex Predictions in CASP16: Are We Making Progress? Proteins 94:106–130.

Zhang Y, Zhang Z, Kagaya Y, Terashi G, Zhao B, Xiong Y, Kihara D (2025a) Distance-AF improves predicted protein structure models by AlphaFold2 with user-specified distance constraints. Communications Biology 8:1392.

Zhang Z, Verburgt J, Kagaya Y, Christoffer C, Kihara D (2025b) Learning with Privileged Knowledge Distillation for Improved Peptide-Protein Docking. ACS Omega 10:26684–26693.

Zhou H, Skolnick J (2011) GOAP: a generalized orientation-dependent, all-atom statistical potential for protein structure prediction. Biophys J 101:2043–2052.

Zhou H, Zhou Y (2002) Distance-scaled, finite ideal-gas reference state improves structure-derived potentials of mean force for structure selection and stability prediction. Protein Science 11:2714–2726.

