## Supplemental Information Doc for "Distance-AF Multimer: A Multi-Step Approach for Protein Complex Structure Modeling with User-Defined Distance Constraints"

**Supplementary Information 1.**

**Phase 2 starting-conformation generation for Distance-AFM**

Algorithm 1 generates geometrically viable starting conformations through a systematic four-stage iterative approach. Native structures are used only for evaluation and do not participate in candidate generation or ranking.

**Algorithm 1:** Distance-AFM Starting-Conformation Generation

***Input:*** *Initial structure X0, restraints D, cluster and intermediate candidate limit K = 3*

***Output:*** *Candidate starting structures Final*

**1:** Candidates ← {(X0, [chain1], [chain2, ..., chainn])}

**2:** Final ← ∅; UsedChainPairs ← ∅

**3:** while Candidates ≠ ∅ do

**4:** Next ← ∅

**5:** for each (X, Done, Remaining) in Candidates, in stored order do

**6:** M ← first(Remaining); Fixed ← all chains except M

**7:** C ← unused restraints between last(Done) and M

**8:** if C = ∅: C ← shortest unused restraint connecting Fixed and M

**9:** if C ≠ ∅: C ← the shortest restraint, or up to three shortest if enabled

**10:** UsedChainPairs ← UsedChainPairs ∪ chain pairs represented in C

**11:** Generated ← ∅

**12:** for each restraint c in C do

**13:** (T, Pivot) ← ComputeTranslationAndPivot(X, c)

**14:** ValidRotations ← ∅

**15:** for each Euler-angle triple sampled at 30° intervals do

**16:** R ← RotationMatrix(angles)

**17:** X′ ← Transform(X, M, T, R, Pivot)

**18:** add R to ValidRotations if X′ passes the clash-count criterion (Cα–Cα < 12 Å)

**19:** Clusters ← connected components of valid grid orientations

**20:** refine every cluster at 15°; add only sterically feasible gap rotations

**21:** Selected ← up to K largest refined clusters (K = 3)

**22:** Representatives ← RotationMatrix(mean Euler angles) for each selected cluster

**23:** append Transform(X, M, T, R, Pivot) for each representative R to Generated

**24:** if Generated = ∅ then Generated ← {X}

**25:** Done′ ← append(Done, M); Remaining′ ← remove-first(Remaining)

**26:** for each Xnew in Generated, in generation order do

**27:** if Remaining′ = ∅ then append Xnew to Final

**28:** else append (Xnew, Done′, Remaining′) to Next

**29:** Candidates ← first K entries of Next

**30:** return Final

**Sequential Chain Incorporation and Candidate Retention (Lines 1–6, 25–29)**

The first chain is the initial reference, and the other chains are processed sequentially. At each assembly stage, the next chain is selected as the moving unit while all other chains remain fixed. Done and Remaining are ordered lists that track processed and unprocessed chains, respectively. Retained candidates are processed in their stored order, and the first K intermediate candidates (K = 3) are retained for the next stage. A two-chain complex requires only one transformation-search stage.

**Transformation Search (Lines 7–18)**

For each candidate, the algorithm first considers unused restraints linking the last processed chain and the selected moving chain. If none exist, it uses the shortest unused restraint connecting the moving chain to any fixed chain. The shortest applicable restraint is used by default, or up to three shortest restraints are searched separately when enabled. Each search translates only the selected moving chain to satisfy its selected restraint, then samples rotations about the fixed-residue pivot at 30° intervals in each of the three Euler angles (α, β, γ). Clashes are identified by inter-chain Cα–Cα distances below 12 Å, and orientations are filtered using the allowed clash-count criterion. Each search initializes its own valid-rotation collection and retains its associated translation and pivot. Used chain pairs are tracked across candidate processing.

**Clustering and Representative Selection (Lines 19–22)**

Valid grid orientations are grouped into connected components. Neighbors differ by one 30° step in exactly one Euler angle, with the other two unchanged. Each cluster is locally refined by exploring angular gaps between neighboring rotations at 15° intervals. Additional rotations are incorporated into the cluster if they satisfy the steric feasibility criteria. After refinement, the K largest clusters (K = 3, or all clusters if fewer are available) are selected by their numbers of valid orientations. For each selected cluster, the mean of each Euler angle (α, β, γ) is calculated over all rotations in the refined cluster. These three mean angles directly define the representative rotation used to generate a starting conformation.

**Candidate Generation and Retention (Lines 23–30)**

The averaged representative rotations are applied to the selected moving chain using the translation and pivot associated with their search, while all other chains remain fixed. The resulting structures are appended in generation order. If no valid transformation is obtained, the current structure is carried forward as the chain lists advance. The first K intermediate candidates are retained without sorting by a combined clash-and-restraint score; terminal-stage structures are saved for Phase 3 fine tuning. Native-reference RMSD, when computed, is used only for evaluation and not for candidate selection.

Overall, Algorithm 1 combines progressive assembly, rigid-body rotational sampling, clustering and local refinement with bounded intermediate candidate retention. It produces geometrically filtered starting conformations for subsequent fine tuning under the user-provided distance restraints.

**Computation time and memory usage**

The computation time for Distance-AFM is largely dependent on the length of the protein sequence and the number of iterations. For a three-chain target complex of up to 500 amino acids total, running 7,000 iterations is generally sufficient. This requires about 4–5 hours on an NVIDIA RTX A6000 GPU with 48 GB of memory. We have optimized the Distance-AF framework to efficiently utilize GPU memory during backpropagation. With an NVIDIA RTX A5500 GPU with 24 GB of memory, it is possible to run any target sequence of up to 3,000 residues.

**Supplementary table legends**

Supplementary Tables 1–8 are provided as separate worksheets in the accompanying Excel file, suppl_table_v1.1.xlsx. The worksheet names below match the file. In Tables 1, 7, and 8, Length is the total number of residues in the modeled complex; chain/domain ranges and constraint residue pairs use concatenated residue numbering. Distances and RMSDs are reported in ångströms (Å). Numerical scores retain the precision available in the source data; displayed values are rounded for readability. Statistical analyses use the underlying unrounded values.

**Supplementary Table 1 (in a separate Excel file). Benchmark protein complexes and distance constraints.**

The 27 benchmark targets comprise 11 two-chain, 8 three-chain, and 8 four-chain complexes. Each row lists the PDB ID, number of chains, total sequence length, chain/domain residue ranges, and the residue pairs and target Cα–Cα distances used as constraints. Up to 12 distance constraints are used per interacting chain pair; fewer are used when fewer residue pairs are successfully matched during constraint generation. Each two-chain target uses 12 constraints, whereas counts vary across chain pairs in the three- and four-chain targets. These targets form the benchmark used in Figures 2–4. Worksheet: “Supp. Table 1. AFM-HCMC”.

**Supplementary Table 2 (in a separate Excel file). Model-quality comparison across the 27-target benchmark.**

Source data for Figure 2a–e and Table 1, with target-level results also used in Figure 3. Each row reports DockQ, fraction of native contacts (Fnat), ligand RMSD (lRMSD), interface RMSD (iRMSD), and RMSD for AlphaFold-Multimer (AFM), Distance-AFM (D-AFM), Protenix, Chai-1, and AlphaLink2 (AL2). DockQ and Fnat are dimensionless and higher values indicate better predictions; lower RMSD values indicate better predictions. Interface scores are averaged over interfaces for multichain targets. “N/A” denotes an unavailable Chai-1 prediction for 6HCP. Worksheet: “Supp. Table 2. Fig. 2a-e”.

**Supplementary Table 3 (in a separate Excel file). Effect of the number of distance constraints.**

Source data for Figure 2f,g. The table contains 88 target–setting combinations: 11 two-chain targets evaluated with 0, 1, 3, 6, 12, 18, 24, or 30 constraints. Columns identify the target, chain count, number and setting of constraints, DockQ, and RMSD. Zero constraints denotes the unconstrained AFM baseline; 12 constraints is the default setting. Worksheet: “Supp. Table 3. Fig. 2f-g”.

**Supplementary Table 4 (in a separate Excel file). Effect of uncertainty in distance constraints.**

Source data for Figure 2h,i. The 11 two-chain targets were evaluated with 12 constraints under exact-distance, ±2.5 Å, and randomized ±5 Å conditions, giving 33 target–condition combinations. Columns specify the condition, range half-width, whether randomization was applied, DockQ, and RMSD. The half-width is zero for exact constraints. Worksheet: “Supp. Table 4. Fig. 2h-i”.

**Supplementary Table 5 (in a separate Excel file). Consensus scoring and structural accuracy of candidate models.**

Source data for Figure 4a–d, comprising 138 candidate models across the 27 benchmark targets. Each row identifies the target, model type (direct D-AFM refinement or a coarse-tuned candidate), chain count, number of models evaluated for that target, Jury score, DockQ, and RMSD. Higher Jury scores indicate stronger consensus support. These model-level data support the global and within-target correlations and top-k overlap analyses in Figure 4. Worksheet: “Supp. Table 5. Fig. 4a-d”.

**Supplementary Table 6 (in a separate Excel file). Component rankings and detailed outputs of the Jury scoring procedure.**

Detailed scoring records for the same 138 models in Supplementary Table 5. The table includes the final Jury score and rank, 23 component ranking inputs, raw scoring metrics, additional Jury RJS outputs, and redundancy-related scores and model identifiers. The 23 inputs combine DFIRE, GOAP, and ITScorePro with 20 criteria from the VoroIF-jury framework. These records document the component scores used in the Figure 4 analysis; raw metrics retain their original names and scales. Worksheet: “Supp. Table 6. Fig. 4 Jury”.

**Supplementary Table 7 (in a separate Excel file). Peptide–protein complexes and distance constraints.**

Target definitions and constraints for the six peptide–protein complexes in Figure 5: 1CN3, 8U51, 8G4Y, 9CK0, 7ZFR, and 5AER. Columns provide PDB ID, chain count, total sequence length, chain/domain residue ranges, and the constrained residue pairs with target Cα–Cα distances. Six constraints were used per peptide, giving 12 constraints for the two-peptide target 5AER and six for each other target. Worksheet: “Supp. Table 7. PBSC”.

**Supplementary Table 8 (in a separate Excel file). Large multimeric assemblies and distance constraints.**

Target definitions and constraints for the five assemblies in Figure 6: 1BHC, 6EN8, 6QVJ, 2KWD, and 4B6J. The modeled systems contain five to ten chains. Each row gives the PDB ID, chain count, total sequence length, chain/domain residue ranges, and all supplied residue-pair distance constraints, with target Cα–Cα distances in Å. Worksheet: “Supp. Table 8. UCAB”.
